# A set of common movements within GPCR-G-protein complexes from variability analysis of cryo-EM datasets

**DOI:** 10.1101/2020.11.27.401265

**Authors:** Jacopo Marino, Gebhard F.X. Schertler

## Abstract

G-protein coupled receptors (GPCRs) are among the most versatile signal transducers in the cell. Once activated, GPCRs sample a large conformational space and couple to G-proteins to initiate distinct signaling pathways. The dynamical behavior of GPCR-G-protein complexes is difficult characterize structurally. Here, we report on the use of variability analysis to characterize the inherent flexibility within the cryo-EM dataset of the rhodopsin-G_i_-protein complex (Tsai et al., 2019), on which this article builds on. We compare the outcome of this analysis with recently published results obtained on the cannabinoid-G_i_- and secretin-G_s_-receptor complexes. Despite differences related to the biochemical compositions of the three samples, a set of consensus movements emerges. We anticipate that systematic variability analysis on GPCR-G-protein complexes may provide useful information not only at the biological level, but also for improving the preparation of more stable samples for cryo-EM single-particle analysis.

## 1. Introduction

G protein-coupled receptors (GPCRs) constitute the most versatile and diverse membrane protein family, with around 800 genes in humans. GPCRs recognize a large variety of extracellular stimuli, and once activated, initiate intracellular signaling cascades to trigger pivotal cellular responses (Sexton, 2018). Those responses include vision, smell, taste, secretion, nerve system regulation, cellular differentiation, and immune response. As consequence, diseases caused by malfunctioning of GPCRs are very diverse and have dramatic outcomes, including cancer. While∼ 30% of FDA-approved medications target GPCRs, an increasing number of lead compounds are on clinical trials (Hauser et al., 2017). However, pharmacological intervention is greatly limited by our understanding of the GPCR signaling pathway, which is complex and only partially understood (Capper and Wacker, 2018; Wacker et al., 2017). There is strong interest in rationalizing the GPCR signally pathway, as this information will undoubtedly translate into better understanding of biased agonism (Rajagopal et al., 2010; Violin et al., 2014) and ultimately to the design of therapeutics with better specificity and less side-effects (Hauser et al., 2017). Within a simplified picture, GPCR signaling is propagated trough trimeric Gαβγ-proteins, where Gα belongs to four major families, G_i/o_, G_s_, G_q/11_ and G_12/13_ (Costa-Neto et al., 2016). Activated GPCRs associate to trimeric G proteins, inducing exchange of GDP to GTP, and dissociation of Gα from Gβγ,triggering intracellular cascades (Wacker et al., 2017). Activated GPCRs are subsequently phosphorylated mostly at their C-terminus by G-protein receptor kinases (Gurevich et al., 2012), and the arrestin signaling pathway follows (Azevedo et al., 2015; Gurevich and Gurevich, 2019; Gurevich et al., 2012). The unlocking of the molecular basis for G-protein selectivity has been initially approached by sequence analysis (Flock 2017). Developments in the biochemistry of GPCRs-G-protein complexes coupled to the selection of stabilizing binders (Qu et al., 2019; Thal et al., 2018), and advances in the field of cryo-electron microscopy (Danev et al., 2020) have resulted in the deposition of more than 300 entries in the Protein Data Bank (PDB), including structures of receptors coupled to long-awaited signaling partners (Huang et al., 2020; Lee et al., 2020; Maeda et al., 2019; Nguyen et al., 2019), and a variety of orthosteric and allosteric ligands (Qu et al., 2019). This growing information on GPCR-G-protein complex structures allows to draw stronger conclusions obtained from structural alignments of GPCRs bound to different classes of G-proteins (García-Nafría and Tate, 2019; Glukhova et al., 2018; Tsai et al., 2019). These analyses reveal the existence of conserved amino-acid contacts between TM3, TM5, and TM6 of GPCRs, and the C-terminal α5 helix of Gα proteins, with some contacts being G-protein subtype specific. However, the interaction between GPCR and G-protein selectivity goes beyond a simple key-and-lock mechanism mediated by the α5 C-terminal helix, and other conserved contacts between GPCRs and G-protein, especially those mediated by intracellular loop 2 (ICL2) and intracellular loop 3 (ICL3) might be essential to ensure proper complex formation (Glukhova et al., 2018; Klenk et al., 2020; Tsai et al., 2019).

The versatility of GPCRs in responding to different ligands, and their capability of coupling to different intracellular partners lies within their highly dynamic nature (Deupi and Kobilka, 2010; Latorraca et al., 2017). Studying the structural dynamics of signaling complexes is a difficult task, and many activated states are transient (Gusach et al., 2020).

Resolving structures of GPCRs, and at the same time retain information about their conformational dynamics, is a difficult task. X-ray free electron lasers (XFELs) have been used to study dynamics of GPCRs (Nass Kovacs et al., 2019; Nogly et al., 2018) as the methodology allows to record movies at very small time scales. However, molecules embedded in the crystal lattice can explore a limited conformational space. Obtaining crystals for GPCR-G-protein complexes is a difficult task due to the amount of material required, but especially due to the conformational heterogeneity naturally present in the sample which will hinder crystal formation.

Within the field of cryogenic electron microscopy (cryo-EM), time-resolved cryo-EM has been employed to resolve conformational states after protein activation (Kontziampasis et al., 2019; Unwin and Fujiyoshi, 2012). Cryo-electron microscopy is suitable for studying conformational heterogeneity (Lyumkis, 2019) because the landscape of conformational states is preserved during sample vitrification (Glaeser, 2018). In standard single-particle analysis, thousands of 2D particles images are used to refine a single, consensus density map to high resolution. However, under these conditions, information about flexibility and conformational heterogeneity is restricted to 3D-classification, within the limitations of the method (Haie-Meder, 1998; Scheres, 2016). Resolving intrinsic structural heterogeneity from 2D particle images is computationally a difficult task, with some approaches being proposed also over the last few years (Dashti et al., 2014; Jin et al., 2014; Tagare et al., 2015; Zhang et al., 2019). Punjani and Fleet have recently included within the cryoSPARC package a new software, namely 3D-variability analysis, based on the Expectation-Maximization algorithm (Punjani and Fleet, 2020). The algorithm allows to explore, within a cryo-EM dataset, particle flexibility and heterogeneity while retaining high-resolution information, including more difficult samples such as small membrane proteins, which is particularly relevant to study structural flexibility of GPCR-G-protein complexes.

At the time of writing of this article, three published 3D-variability analysis performed on GPCR-G-protein have been reported. Here, we compare the outcome of these analysis, and report on the results obtained by performing the same analysis on the rhodopsin-G_i_αβγ-Fab16 complex dataset (Tsai et al., 2019). Interestingly, a set of common movements is shared among the receptors analyzed.

## 2. Materials and Methods

A particle stack of 111357 images from the consensus refinement performed in Relion (Tsai et al., 2019) was imported into cryoSPARC (Punjani et al., 2017) using pyem (Asarnow, D., Palovcak, E., Cheng, 2019). An *ab-initio* map was produced, which was used as input for refinement using a mask that includes flexible regions (HA-domain of G_i_α, and part of G_i_α below TM5-TM6 of rhodopsin) that were previously excluded to achieve higher resolution (Tsai et al., 2019). For 3D-variability analysis, a mask that excludes the detergent micelle was created (**Supplementary Figure 1A**), as previously described (Punjani and Fleet, 2020). For consistency with the other two published analysis with which comparisons are made (Dong et al., 2020; Punjani and Fleet, 2020), three modes were indicated, and the output contained for each mode 20 density maps, which were used for creating movies in Chimera (Pettersen et al., 2004). The first and last frames (frame 0 and frame 19), corresponding to negative and positive values of the reaction coordinate for each variability component (Punjani and Fleet, 2020) were used for displaying differences within each component.

## 3. Results

### 3.1 3D-variability analysis of the rhodopsin-G_i_-Fab16 complex

The results of the 3D-variability analysis obtained on the rhodopsin-G_i_βγ-Fab16 dataset follow. The first component resolves a large movement of the G_i_α alpha-helical (HA) domain, a folding unit characteristic of Gα proteins (Joseph P. Neol, 1993), which due to its flexible connection to the Ras domain remains invisible in most structures. As shown in **Figure 1**, within the 3D reconstructions of the variability component one, the HA-domain is strongly visible in frame zero (blue) where it associates to Gβ, but then becomes invisible in frame 19 (yellow), where it moves away from Gβ (**Figure 1A, and Suppl. Video 1**), thus resembling fluctuations between an open- and closed-conformation observed previously in computational studies (Dror et al., 2015; Sun et al., 2018). We observed the presence of the HA-domain within the rhodopsin-G-protein complex during 3D classification in Relion in one of the four 3D-classes obtained (Tsai et al., 2019). Interestingly, in our analysis, the appearance and disappearance of the HA domain is associated to twisting of rhodopsin around an axis parallel to the membrane, with larger movements especially visible for transmembrane (TM) segment 4 and TM6 (**Figure 1B and C, Suppl. Movie 1**).

**Figure 1.**
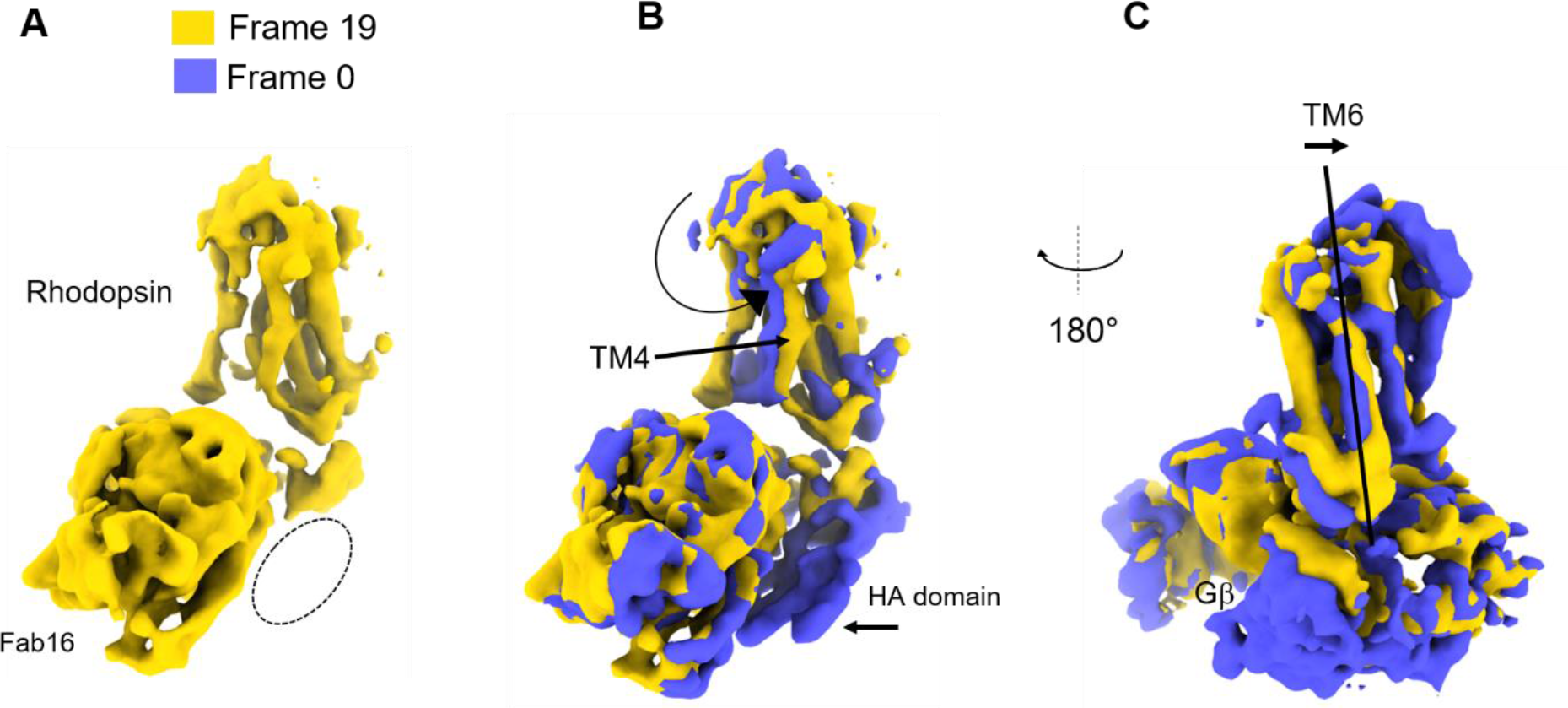
Result component 1 of cryoSPARC 3D variability analysis of the rhodopsin-G_i_-complex bound to Fab16. A) Density map corresponding to frame 19 of the output of component one. In circle is indicated the missing density of the alpha-helical (HA) domain, which is instead visible in frame zero (B). Component one also resolves rotation of rhodopsin along the membrane axis, with larger displacement of TM4 (B) and TM6 (C).

The second component of the variability analysis resolves bending of rhodopsin ∼20° away from Gβ along the axis perpendicular to the membrane, and Fab16 and the receptor oscillates against each other (**Figure 2A)**. Interestingly, when rhodopsin bends towards Gβγ (blue), intracellular loop 2 (ICL2) contacts the N-terminal helix of G_i_α, an interaction absent when rhodopsin is bent away (yellow) **(Figure 2B)**. The interaction of ICL2 with the N-terminal helix of G_i_α is a recurrent feature observed in structures of GPCR-G-protein complexes, and it might be needed to stabilize the receptor-G-protein interface (García-Nafría and Tate, 2019; Glukhova et al., 2018; Tsai et al., 2019). Bending of rhodopsin away from Gβγ produces the effect of allowing TM5 and TM6 to get closer to G_i_α **(Figure 2C)**, thus extending the interaction of ICL3 with G_i_α. The interaction of ICL3 with G_i_α is also a feature that has been already observed in other GPCR-G-protein complexes (García-Nafría and Tate, 2019; Glukhova et al., 2018).

**Figure 2.**
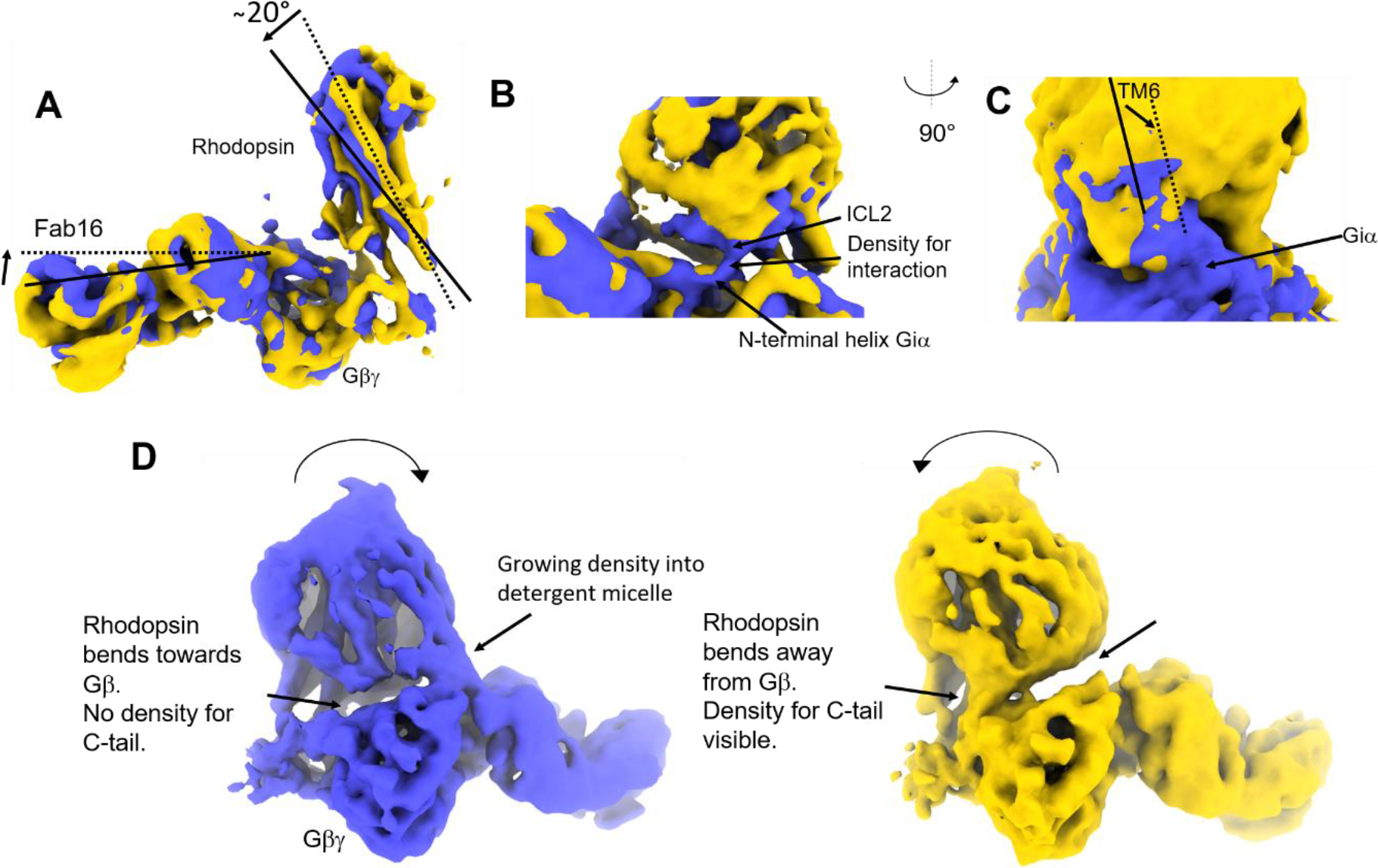
Result component two. A) Representation of the two density maps corresponding to frame zero (blue) and frame 19 (yellow) resulting from component two. The oscillating movement of rhodopsin towards or away from Gβγ is indicated. In B) is shown the density growing from the N-terminal helix of G_i_α, which is not visible in frame 19 (yellow). In C), rotation of rhodopsin causes TM6/ICL3 to make a larger contact area with G_i_α· D) Indicated is the absence (blue) or presence (yellow) corresponding to the rhodopsin C-terminal tail (see also Suppl. Figure 2), and presence (blue) or absence (yellow) of density which might be attributed to posttranslational modifications of Gβγ·

Interestingly, bending of the receptor towards Gβγ allows to visualize the density corresponding to the rhodopsin C-terminal tail (C-tail), which in our rhodopsin-complex (Tsai et al., 2019) and in the muscarinic receptor complex (Maeda et al., 2019) is bound to Gβ. However, our analysis shows that the interaction of the C-tail to Gβ might be dynamic. The density corresponding to rhodopsin’ C-tail is not visible when the receptor bends away from Gβ (blue), also at very low thresholds **(Figure 2D and Suppl. Fig. 2**), indicating that rhodopsin must be at a closer distance to Gβ to stabilize the C-tail in that position.

Another observation that can be made within these reconstructions is that in frame zero of component two, a density growing from Gβ to the receptor micelle is visible **(Figure 2D)**. This density may be attributed to posttranslational modifications of Gβγ interacting with the detergent micelle (Vögler et al., 2008), as Gβγ was purified from bovine retina and thus posttranslational modifications should be retained (Tsai et al., 2019). This density is not present when the receptor bends away from Gβγ (Figure 2D, yellow), also at lower thresholds **(Suppl. Movie 2)**.

Component three resolves transversal bending of rhodopsin, Fab16 and Gβγ. Differently from the movements observed in component two, rhodopsin does not bend towards Gβγ but rather towards the N-terminal helix of G_i_α. Here, Gβγ and Fab16 rotate of ∼ 15° (**Figure 3A**). The rotation of Fab16 propagates on its anchoring points, which are Gβ and the N-terminal helix of G_i_α. The N-terminal helix of G_i_α thus translates vertically, as indicated in **Figure 3B**, thus pulling rhodopsin up and down. The relative video is provided as **Suppl. Movie 3**.

**Figure 3.**
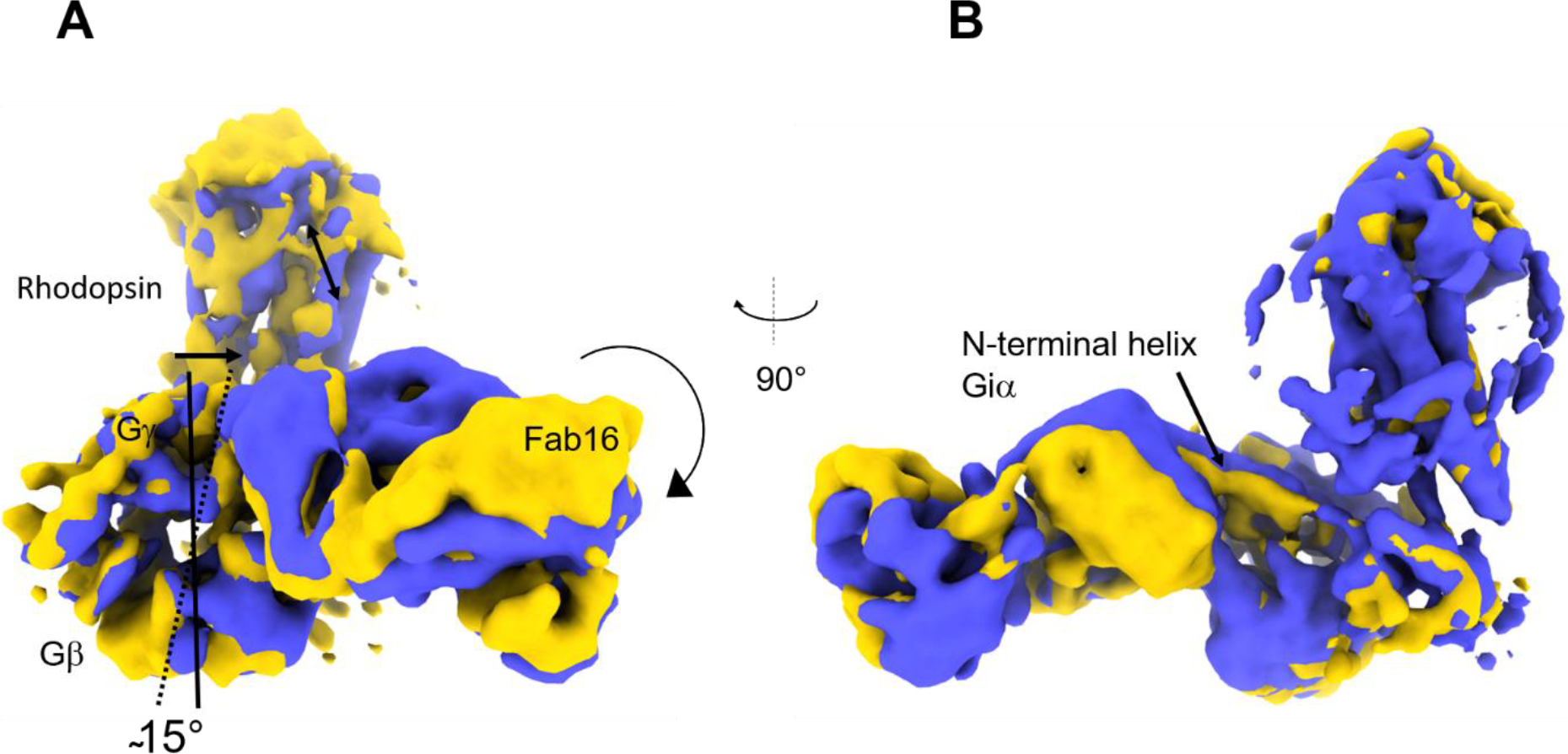
Result component three. A) Representation of the two density maps corresponding to frame zero (blue) and frame 19 (yellow) resulting from component three. A) Rotation of Gβ and Fab16 is indicated. Rotation of Fab16 pulls down the N-terminal helix of G_i_α as indicated in B), causing also rhodopsin to witness a transversal movement of similar amplitude.

### 3.2 Comparison with reported 3D-variability analysis of GPCR complexes

The movements observed within the rhodopsin-Gi-Fab16 complex can be summarized as it follows. The first component resolves twisting of rhodopsin along an axis parallel to the membrane, accompanied by a large movement of the HA-domain. The second component resolves bending of the receptor against Gβγ,and the forming and breaking of interactions at the receptor-G-protein interface; the third and last component resolves a rotation of Fab16 and Gβγ, which propagates to transversal bending of rhodopsin (**Figure 4A**).

**Figure 4.**
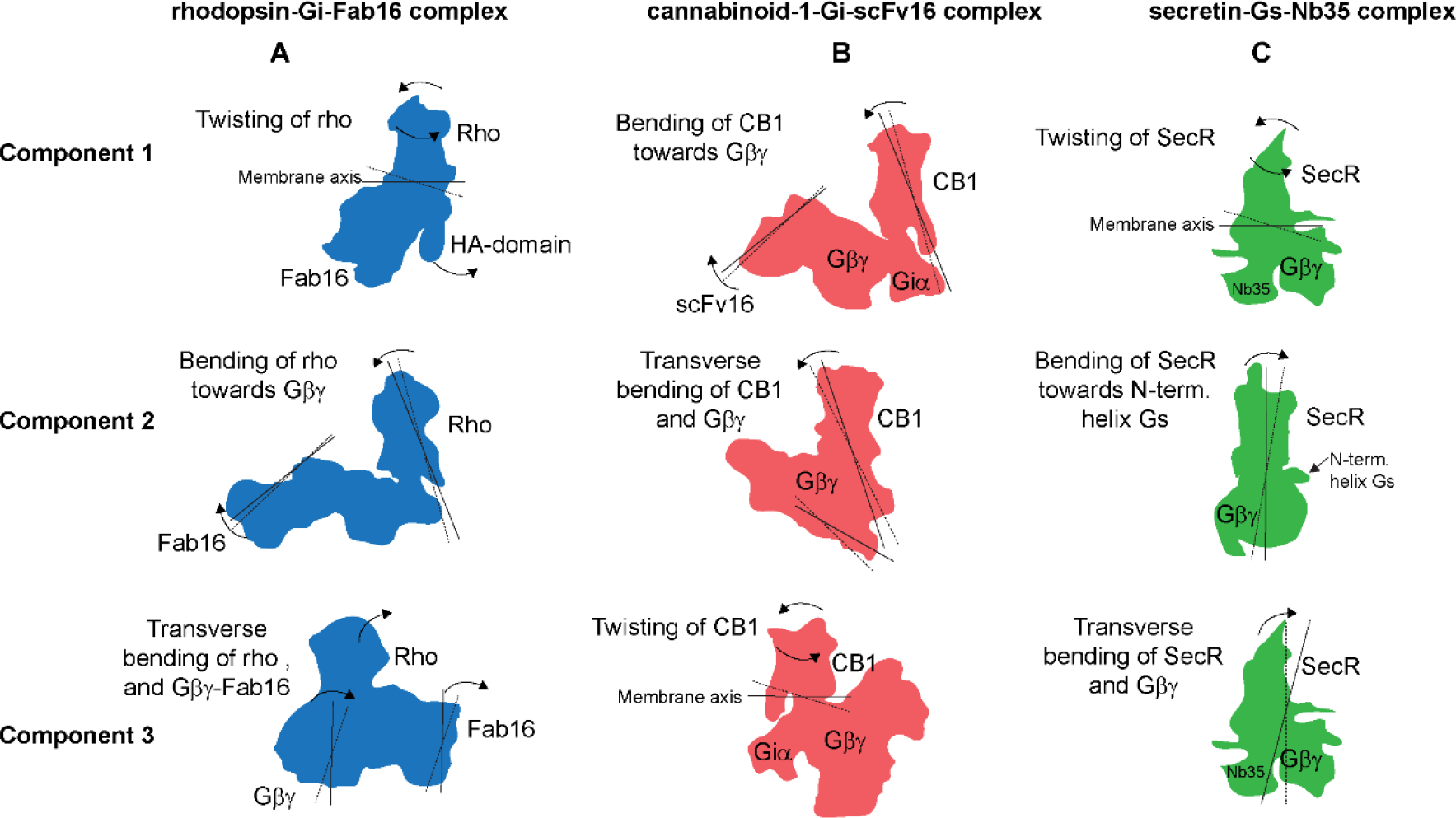
A) Schematic representation of the 3D-variability analysis results for the rhodopsin-G_i_βγ-Fab16 complex. In B) a summary of the results published for the cannabinoid-1-G_i_βγ-scFv16 complex (Punjani and Fleet, 2020), and in C) for the secretin-G_s_βγ-Nb35 complex (Dong et al., 2020). Videos created from the variability analysis can be viewed [here](https://www.notion.so/3D-Variability-Analysis-3DVA-5f351d70f18042e6b32c215372639335) for the cannabinoid receptor, and [here](https://www.notion.so/3D-Variability-Analysis-3DVA-5f351d70f18042e6b32c215372639335) for SecR, while videos for the rhodopsin complex are provided in the Supplementary Materials.

Punjani and Fleet have applied 3D-variability analysis to the cannabinoid receptor 1-G_i_-protein complex dataset (Krishna Kumar et al., 2019) (EMPIAR: 10288). Videos from their analysis can be viewed [here](https://www.notion.so/3D-Variability-Analysis-3DVA-5f351d70f18042e6b32c215372639335). In their study, the analysis in restricted to three components, and in **Figure 4B** we report a schematic representation of the moments that the authors describe (Punjani and Fleet, 2020). The first component shows that the cannabinoid receptor bends towards Gβγ; the second component resolves a motion of Gβγ with transversal bending of the receptor, while the third component resolves twisting of the receptor parallel to the membrane axis (**Figure 4B**).

In a recent study, Dong and colleagues performed 3D variability analysis on the human secretin receptor coupled to G_s_ (Dong et al., 2020). Their analysis takes into consideration three components, and they show very similar movements as compared to the cannabinoid receptor complex. The first component resolves rotation of the receptor along the axis parallel to the membrane. The second component resolves bending of the receptor towards the N-terminal helix of G_s_ (or towards Nb35), a movement with similar amplitude to what observed in component 1 of the cannabinoid receptor analysis, but with a ∼45° twist along the membrane axis. The third component resolves transverse bending of the receptor (**Figure 4C**). Recently, 3D-variability analysis was performed on the Adrenomedullin receptors AM1 and AM2 datasets (Liang et al., 2020). Here, large motions were observed especially within the extracellular domain (ECD), which is a prominent feature of this receptor family. While in the reported analysis the HA-domain of the respective G-proteins is not visible or visible only partially in the cannabinoid-1 reconstruction, more interestingly, the overall motions observed for the cannabinoid-1 and secretin receptor complexes are very similar to what we report for the rhodopsin-complex **(Figure 4)**.

## 4. Discussion

Proteins adopt different conformations along the energy landscape to perform their biological function. Protein flexibility, including disorder, is key to explain molecular machines at work. The complex signal transduction pathway across the cell membrane initiated by G-protein-coupled receptors (GPCRs) is a showcase for transient protein interactions, and structural pharmacology would benefit from characterizing the molecular basis for the transient interactions taking place after GPCR activation. This information will undoubtedly contribute to rationalizing G-protein selectivity and biased agonism (Sexton and Christopoulos, 2018), translating into the development of more specific treatments (Hauser et al., 2017).

Transient interactions between GPCR and G-proteins are difficult to capture by standard structural biology approaches. These events, among others, include G-protein pre-coupling and selectivity, and short-living GDP-bound states (Gusach et al., 2020). The study of dynamics of activated GPCRs has been approached with different techniques including NMR, EPR, FRET, and X-FEL (Gusach et al., 2020). Within true cellular membranes, valuable insights have been produced by single-molecule imaging of GPCR complexes (Sungkaworn et al., 2017).

Cryo-electron microscopy has streamlined many steps to solve atomic-resolution structures of GPCR-G-protein complexes (Danev et al., 2020), and an advantage about cryo-EM is that information about inherent flexibility and sample heterogeneity is preserved within cryo-embedded particles. New algorithms to extract information about dynamics within cryo-EM datasets are providing useful insights about motions connected to biological functions (Dashti et al., 2020; Frank and Ourmazd, 2016; Nešić et al., 2020; Ramírez et al., 2019; Rogala et al., 2019; Stanishneva-Konovalova et al., 2020; Wrapp et al., 2020), including for GPCR-G-protein complexes (Dong et al., 2020; Liang et al., 2020; Punjani and Fleet, 2020).

Here, we report on the outcome of variability analysis performed on the rhodopsin-G_i_-Fab16 dataset (Tsai et al., 2019), and compared the results to the variability analysis performed on the cannabinoid-1 (Punjani and Fleet, 2020) and secretin-receptor (Dong et al., 2020) -G-protein complexes.

The three GPCR-G-protein complexes compared present similarities and differences. Cannabinoid and rhodopsin are class A GPCRs, are bound to G_i_, have the same binder (scFv16 and full Fab16, respectively), and all three complexes are embedded within a LMNG micelle. But they also have important differences. The secretin receptor is a class B GPCR bound to G_s_ and has a nanobody (nb35) bound. However, upon analyzing the outcome of the variability analysis, these complexes share remarkable similarities in terms of the movements observed, and these movements be categorized within three sets: 1) rotation of the receptor on an axis parallel to the membrane; 2) bending of the receptor towards Gβγ (cannabinoid and rhodopsin) or towards the N-terminal helix of G_s_ (secretin) on an axis perpendicular to the membrane; 3) transversal bending of the receptor relative to Gβγ·.

The results obtained provide insights into dynamical features of GPCR-G-protein complexes that are not always captured in high-resolutions structures, and which might relate to biological and biochemical aspects of complex formation. One of these features is the HA-domain of Gα, which was observed to fluctuate from a Gβ-bound state to a closed-state in molecular dynamics simulations (Dror et al., 2015; Sun et al., 2018) and was reported only in a few structures (Kang et al., 2018; Klenk et al., 2020; Rasmussen et al., 2011). In these structures, the position of the HA-domain in its open conformation, very similar to what we observe within the frames of component one of our analysis (**Figure 1**), within a motion that is propagated to the Ras domain. These movements are in synchrony with rotation of rhodopsin on an axis parallel to the membrane. The large movements of the HA-domain might recapitulate similar conformational changes occurring during GDP release, as separation of the HA-domain from the Ras domain might contribute to increase the probability of nucleotide release (Dror et al., 2015; Sun et al., 2018).

Another aspect of GPCR-G-protein complexes related to flexibility concerns the interactions at the interface between receptor and G-proteins, especially those interactions mediated by the receptors’ intracellular loops (ICL). Structural alignments indicate that ICL2 and ICL3 often, but not always, are at contact distance to Gα (García-Nafría and Tate, 2019; Glukhova et al., 2018). In the structure of the NTR1 receptor-G_i_ complex in the canonical (C) state, ICL2 and ICL3 make a larger number of contacts to Gα as compared to the structure in the non-canonical (NC) state, which is rotated as compared to the C-state (Klenk et al., 2020). In our analysis, the interactions between rhodopsin ICL2 and ICL3 and Gαβ are breaking and forming along the oscillations and rotations observed, and might recapitulate a transition between two different states not far from what observed in the NRT1 structures (Klenk et al., 2020), albeit within a smaller degree of conformational flexibility.

Concluding, the characterization of intrinsic protein flexibility and heterogeneity embedded within cryo-EM datasets is becoming a routine analysis performed on the side of the traditional single-particle analysis. Together with data obtained with other methods to study GPCRs (Gusach et al., 2020), this information will contribute to rationalize the complex and dynamic nature of the GPCR signaling pathway, and can be useful to select specific binders that limit the degree of freedom between receptor and G-proteins to facilitate high-resolution structure determination by cryo-EM.

## 5. Acknowledgments

Jacopo Marino and Gebhard F.X. Schertler are thankful to the colleagues who contributed to the original eLife publication on the rhodopsin-G-protein complex, and to especially Ricardo Adaixo, Filip Pamula, Jonas Mühle, and Ching-Ju Tsai.

## 6. Author contributions

Jacopo Marino conceived the project with Gebhard F.X. Schertler. Jacopo analyzed data and wrote the manuscript. Gebhard F.X. Schertler helped writing the manuscript.

## 7. Competing interests

Jacopo Marino declares no competing interests. Gebhard Schertler declares that he is a co-founder and scientific advisor of the company leadXpro AG and InterAx Biotech AG.

## 8. Funding

Jacopo Marino acknowledges a Swiss National Science Foundation (SNSF) grant #19082). Gebhard F.X. Schertler acknowledges SNSF grants # 310030B_173335 and # 310030_192760.

## Supplementary Material

**Supplementary Figure 1.**
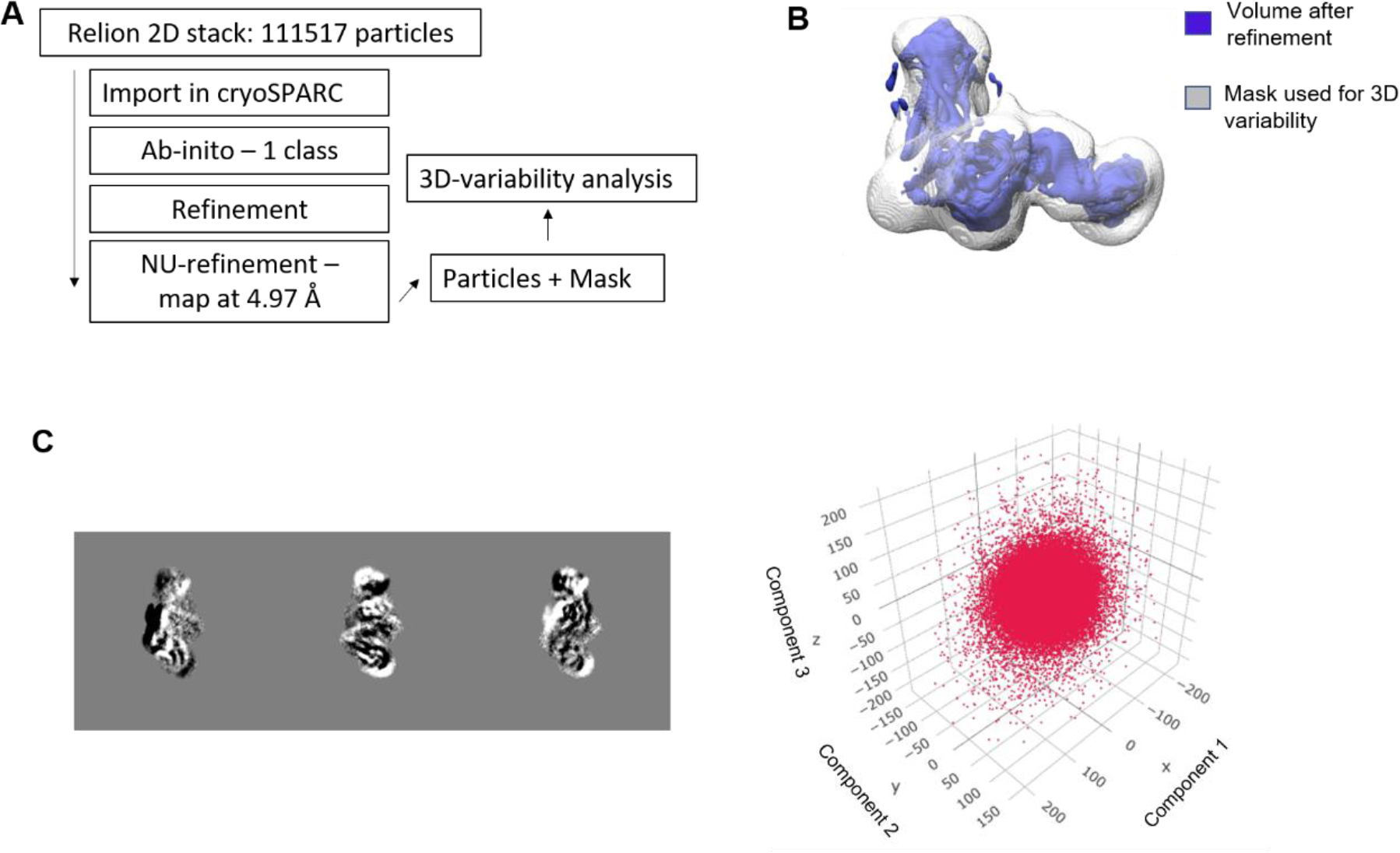
A) Processing pipeline in cryoSPARC. B) Mask (volume in blue) created for the 3D-variability analysis. The detergent micelle, in red, was excluded from the analysis. C) Output from the 3D-variability display of cryoSPARC.

**Supplementary Figure 2.**
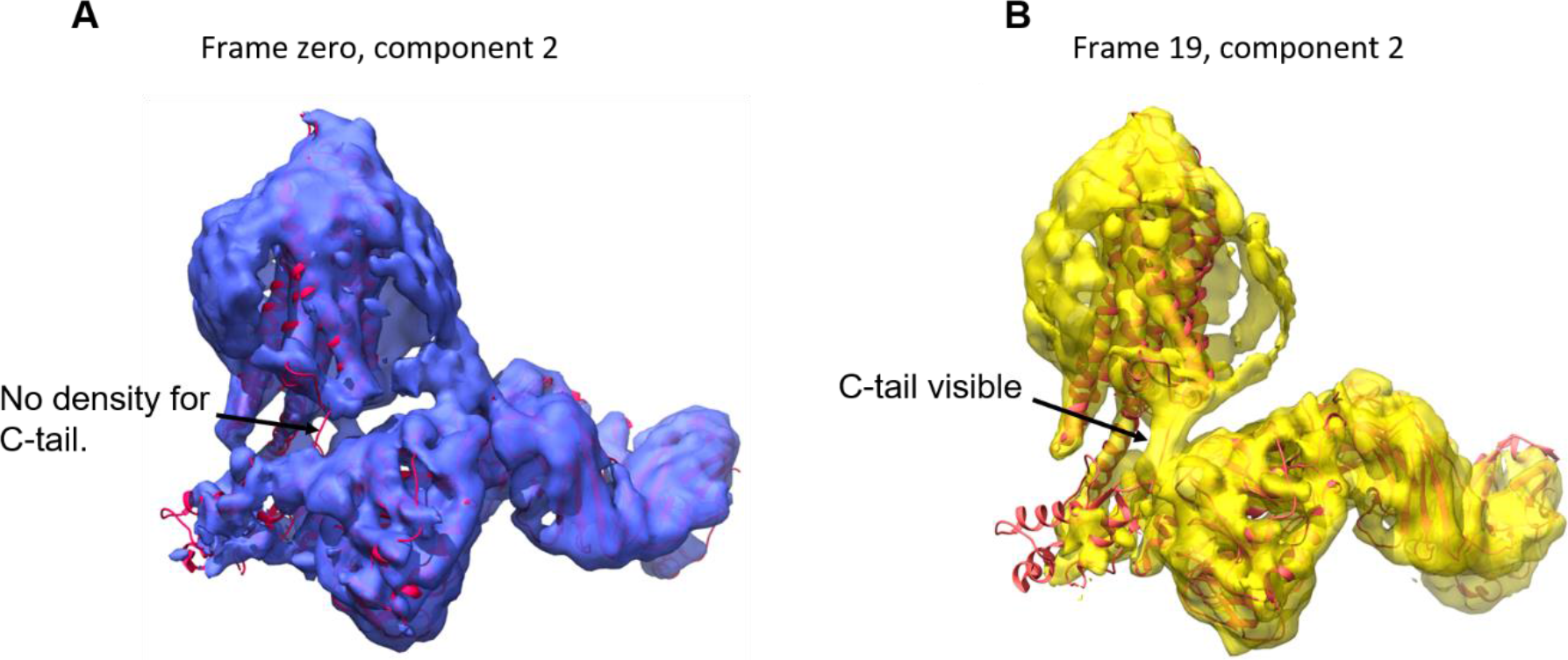
Fit of the atomic model of the rhodopsin-G-protein complex (PDB:6QNO, in red) into density maps for frame zero (A), and frame 19 (B), obtained within component 2 of the 3D-variability analysis.

**Supplementary Video 1**

**Supplementary Video 2**

**Supplementary Video 3**

## Notes

### Competing Interest Statement

The authors have declared no competing interest.

